# 3D bioprinting directly affects proteomic signature and myogenic maturation in muscle pericytes-derived human myo-substitute

**DOI:** 10.1101/2023.07.10.548389

**Authors:** Alessio Reggio, Claudia Fuoco, Francesca De Paolis, Stefano Testa, Nehar Celikkin, Sergio Bernardini, Jacopo Baldi, Roberto Biagini, Dror Seliktar, Carmine Cirillo, Paolo Grumati, Stefano Cannata, Marco Costantini, Cesare Gargioli

## Abstract

Skeletal muscle tissue engineering (SMTE) has recently emerged to address major clinical challenges such as volumetric muscle loss. Here, we report a rotary wet-spinning (RoWS) biofabrication technique for producing human myo-substitutes with biomimetic architectures and functions. We show how the proposed technique may be used to establish a well-tailored, anisotropic microenvironment that promotes exceptional myogenic differentiation of human skeletal muscle-derived pericytes (hPeri). Using high-resolution mass spectrometry-based proteomics with the integration of literature-derived signaling networks, we uncovered that i) 3D biomimetic matrix environment (PEG-Fibrinogen) confers a lower mitogenicity microenvironment compared to standard 2D cultures, favoring the formation of contractile-competent bundles of pericytes-derived myotubes in an anchoring-independent 3D state, and ii) the bioprinting method promotes an upregulation of muscle matrix structural protein besides increasing contractile machinery proteins with respect to 3D bulk cultures. Finally, *in vivo* investigations demonstrate that the 3D bioprinted myo-substitute is fully compatible with the host ablated muscular tissue, exhibiting myo-substitute engraftment and muscle regeneration in a mouse VML model. Overall, the results show that 3D bioprinting has a superior capability for controlling the myogenic differentiation process on a macroscale and, with future refining, may have the potential to be translated into clinical practice.

## 1. Introduction

Over the last decade, biofabrication technologies have brought a new wave of enthusiasm in the tissue engineering community as a potential option for the fabrication of architecturally complex and functionally biomimetic tissue/organ substitutes. The increasing interest in such systems is justified by their unprecedented advantages that enable building multi-material, cellularized constructs with high spatial resolution, repeatability and 3D biomimetic structural complexity.

In the context of skeletal muscle tissue engineering (SMTE), biofabrication technologies have demonstrated great potential in guiding the alignment and differentiation of myogenic precursors – including human primary cells – both in *in vitro* and *in vivo* models. ^1–4^. Notable examples in the field include the assembly of multi-cellular models for the *in vitro* recapitulation of the neuromuscular (NMJ) or myotendinous (MTJ) junction complexity, mimicking exercise and pharmacological response, or the fabrication of artificial skeletal muscles for regenerative medicine applications in small animal models ^5–8^. Moreover, biofabrication strategies have been proposed for the manufacturing – though to a small scale (approx. 1 cm^3^) – of whole cut, bovine meat-like tissue containing muscle, fat, and vasculature tissue ^9,10^.

Despite such a great and encouraging results, various biological fundamental aspects remain to be elucidated. In particular, it has not been investigated thoroughly how the bioprinted microenvironment rewires muscle progenitors to form functional muscle bundles. Moreover, it is not clear whether there exists a difference in terms of myogenic process modulation between biofabricated muscles and engineered muscle constructs obtained using conventional approaches (e.g. 2D or bulk hydrogels). Such missing information is of the utmost importance as it may help to unravel the complexity of neo-muscle tissue formation, and, as a consequence, to develop improved strategies for its assembly *in vitro*.

Herein, a new biofabrication strategies – namely 3D rotary wet-spinning (ROWS) – supported by an innovative microfluidic printing head was employed to fabricate biomimetic, core/shell fiber hydrogel constructs where human skeletal muscle-derived pericytes (hPeri) were encapsulated. As reported in previous studies, skeletal muscle derived hPeri are a powerful source of competent myogenic precursor cells and such myogenic potential is favored by culturing these cells in three-dimensional (3D) hydrogel matrices^11^. The myogenic capacity of the biofabricated hPeri-loaded samples was first evaluated *in vitro,* and benchmarked against pericytes grown in 2D or within 3D bulk hydrogels. To pinpoint molecular details at proteome resolution level among the three experimental groups, we exploited state-of-the-mass spectrometry-based proteomics to dissect molecular information from pericyte-based myo-substitutes. Notably, the proteomic analysis revealed that the 3D environment promotes lower mitogenicity levels together with downregulation of adhesion molecular pathways respect to 2D cultures, favoring the formation of contractile-competent bundles of pericytes-derived myotubes. Moreover, the biofabricated myo-substitutes showed a significant upregulation of both muscular matrix structural and contractile machinery proteins with respect to 3D bulk cultures. Finally, to demonstrate the robustness of the proposed strategy, biofabricated myo-substitutes were implanted into a mouse model of muscle volumetric damage to assess their grafting and integration capabilities.

Altogether, our results highlight that biofabrication strategies may represent the next golden-standard for the recapitulation of the skeletal muscle architecture, and, more generally, for the assembly of functional artificial muscle substitute.

## 2. Results and Discussion

### 2.1 Microfluidic-assisted 3D rotary wet-spinning bioprinting

To produce highly aligned muscle fibers from myogenic competent human pericytes (hPeri), we developed and validated a new biofabrication platform to reproduce such complex architectural organization. The system leverages the principles of wet-spinning technologies for the continuous, volumetric production of hydrogel fibers (Figure 1A). Contrary to conventional extrusion systems, our bioprinter is composed of just two axes: one linear (Figure 1A, *x-axis* arm) for the translational movement of the printing head, and one rotary for the collection of the wet-spun fibers. A core part of the whole system is represented by the wet-spinning microfluidic printing head (ws-MPH). Specifically, we designed such microfluidic device to enable the continuous production of core/shell hydrogel fibers (Figure 1B). To this aim, the MPH is equipped with a co-axial extrusion nozzle which is placed at the bottom of a crosslinking bath microtank. Core and shell hydrogel precursor solutions are supplied respectively to the inner and outer nozzle of the co-axial system through a network of microchannels embedded in the microfluidic device. This configuration enables the formation of a stable co-axial flow of the two solutions within the nozzle which, upon extrusion in the bath microtank, undergoes instantaneous gelation resulting in a core/shell hydrogel fiber. Notably, the direct mounting of a crosslinking bath microtank over the ws-MPH not only significantly reduces the overall dimension of the wet-spinning printing head, but also greatly simplify the production and deposition of core/shell fibers as it removes the need of using multi-axial nozzles and the supplying of a third solution to crosslink the core/shell fiber.

**Figure 1.**
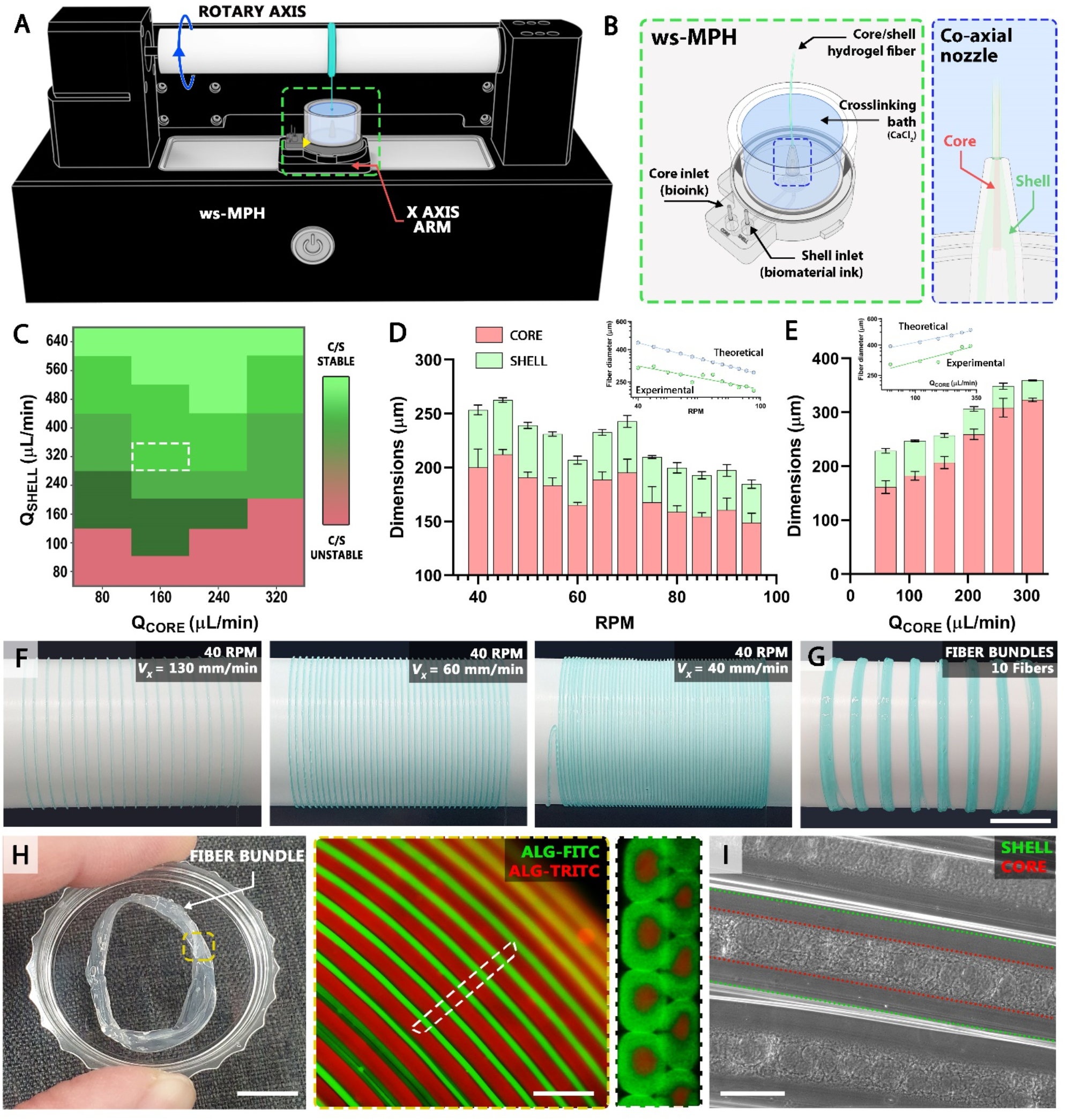
Microfluidic-assisted 3D rotary wet-spinning (RoWS) bioprinting. A) Schematic representation of the 3D RoWS bioprinting platform and B) the co-axial microfluidic printing head (MPH) used to manufacture core/shell hydrogel fibers. C) Stability diagram displaying the Q_SHELL_ and Q_CORE_ values suitable for continuous fiber spinning. D) Influence of rotational speed over the size of extruded fibers (Q_SHELL_ = 320 μL/min, Q_CORE_ = 160 μL/min). E) Effect of different Q_CORE_ over core/shell dimensions, keeping constant Q_SHELL_ (320 μL/min) and the rotational speed (40 rpm). Insets in D, E) highlight the difference between theoretical and experimental values of fiber sizes, most likely caused by alginate gel shrinking upon crosslinking. F) Various spiral fiber patterns obtained for different translation speeds; G) assembly of fiber bundles with a specific number of fibers. H, I) Bright field and fluorescent images of highly aligned fiber bundles characterized by a core/shell architecture. Scale Bars: h) (left) 5 mm, (right) 500 μm, i) 250 μm.

Before starting the experiments with cells, we explored the full potential of our system by investigating the operational ranges where a stable production of core/shell fibers can be achieved. To this aim, we studied the influence of three main process parameters: the shell and core ink flow rates (Q_SHELL_, Q_CORE_) and the rotational speed used for fiber collection. During this preliminary characterization, the composition of the two inks was not varied. Respectively, the core bioink contained a semi-synthetic biopolymer – namely PEG-Fibrinogen (PF), at a concentration of 8 mg/mL – extensively used in skeletal muscle tissue engineering, while the shell biomaterial ink was composed of a mixture of high and low molecular weight alginates – at a respective concentration of 10 mg/mL and 20 mg/mL – to enable the immediate gelation with the Ca^2+^ ions contained in the microtank. The results of our preliminary system characterization are shown in Figure 1C-E. We first run experiments at a constant rotational speed of 40 rpm, to scan the effects of ink flow rates over fiber extrusion stability. A stable production of core/shell fibers can be achieved for a broad set of ink flow rates when the Q_SHELL_ is higher than 160 μL/min, while below such values, the fiber extrusion process resulted unstable, with frequent break-up of the extruded fibers. Accordingly, we selected as suitable values for the ink flow rates the pair Q_SHELL_ = 320 μL/min and Q_CORE_ = 160 μL/min. These values represent a proper trade-off between the system throughput and stability, and cell friendliness, minimizing the potential shear stress that cells could experience during extrusion at higher flow rates. Following these experiments, we characterized the influence of rotational speed over core and shell sizes using the selected pair of ink flow rates. The results revealed an opposite relationship, with faster rotational speeds causing the production of smaller fibers (Figure 1D). This can be explained by the fact that fibers collected on a drum that rotates more quickly underwent a higher level of stretching. Additionally, we characterized the effect of different Q_CORE_ over core/shell dimensions, keeping constant Q_SHELL_ (320 μL/min) and the rotational speed (40 rpm). As shown in Figure 1E, an exclusive increase of Q_CORE_ results in fibers having larger cores and thinner shells. Interestingly, it was consistently reported that experimental values were lower than theoretical ones. Such effect seemed unrelated to rotational speed (Figure 1D, E) where the experimental and theoretical data are plotted in a log-log space forming two parallel lines, and it was likely caused by gel shrinking effects that occur during the crosslinking of alginate solutions.

After fiber extrusion characterization, we proceeded with testing the capacity of our rotary wet-spinning bioprinter in terms of fiber spatial deposition precision. Notably, the developed custom user interface enables to independently control the X-axis motion and the rotational speed, allowing the creation of less or more densely packed spiral patterns of fibers that can be used to form thicker, rod-shape 3D fiber constructs (Figure 1F). In this study, however, we focused on the production in series of bundles composed of highly-anisotropic, compartmentalized hydrogels composed of core/shell fibers (Figure 1G) as these closely resemble skeletal muscle architecture. As shown in Figure 1H, the fabricated bundles enjoy a high mechanical stability that allows for an easy removal from the drum. Fibers within the bundles result tightly packed and aligned with a distinct and precise core and shell spatial compartmentalization. Of note, the latter can be also observed using phase contrast light when the core and shell hydrogels have either different chemical composition (i.e. different optical densities) or different microstructure as in the case of fibrous fibrin core and smooth alginate shell (Figure 1I).

### 2.2 From 2D to 3D bioprinting: promoting *in vitro* human pericytes myogenic structuration

To effectively and quantitatively evaluate the capacity of the proposed bioprinting system in guiding on a large scale (i.e. centimeter scale) the myogenic differentiation of freshly harvested skeletal muscle-derived human pericytes (hPeri), we defined and benchmarked three experimental groups: i) a 2D standard cell culture system, ii) a 3D soft, bulk hydrogel and iii) 3D bioprinted constructs. The first two groups have been selected as they represent distinct culturing conditions largely employed and characterized among the skeletal muscle research community ^12, 13^, thus being proper references to investigate whether a faithful muscle architecture can be obtained using our own biofabrication strategy.

Thus, we run in parallel a series of experiments where hPeri structural organization and myogenic differentiation were first assessed using phase contrast and confocal microscopy. For all 3D experiments (i.e. bulk and core phase in bioprinted fibers), we used as hydrogel matrix photocurable PEGylated-Fibrinogen (PF), a semi-synthetic biopolymers that has shown in the past great capacity in supporting *in vitro* the differentiation of skeletal muscle precursors ^2, 3, 12, 14, 15^. The results are shown in Figure 2. As it can be easily appreciated in Figure 2A-F, hPeri showed a pronounced differentiation in all the investigated groups, with the formation of multi-nucleated myotubes highly positive for myosin heavy chain (MyHC, a key muscle marker demonstrating advanced myogenic differentiation). However, the structural and spatial organization of these myotubes were significant different among the 3 groups. In the 2D cell culture group, hPeri differentiated into a single layer of adjacent myotubes exhibiting a short-range, random spatial alignment (Figure 2A, B). In the 3D bulk hydrogel group, hPeri formed an entangled network of myotubes with an overall isotropic spatial distribution (Figure 2C-2E). In the 3D printed group, hPeri were viable (Supplementary FigureS1) and developed into densely packed myotubes characterized by a unidirectional, long-range alignment (Figure 2F). During differentiation, biofabricated constructs showed spontaneous, synchronous contraction of large samples area, denoting a high level of maturation (Supporting Video 1). Remarkably, spontaneous *in vitro* contraction in human derived myogenic progenitors is generally hard to attain, unless upon mechanical or electrical stimuli, thus providing the robust demonstration of myo-structure thoroughness.

**Figure 2.**
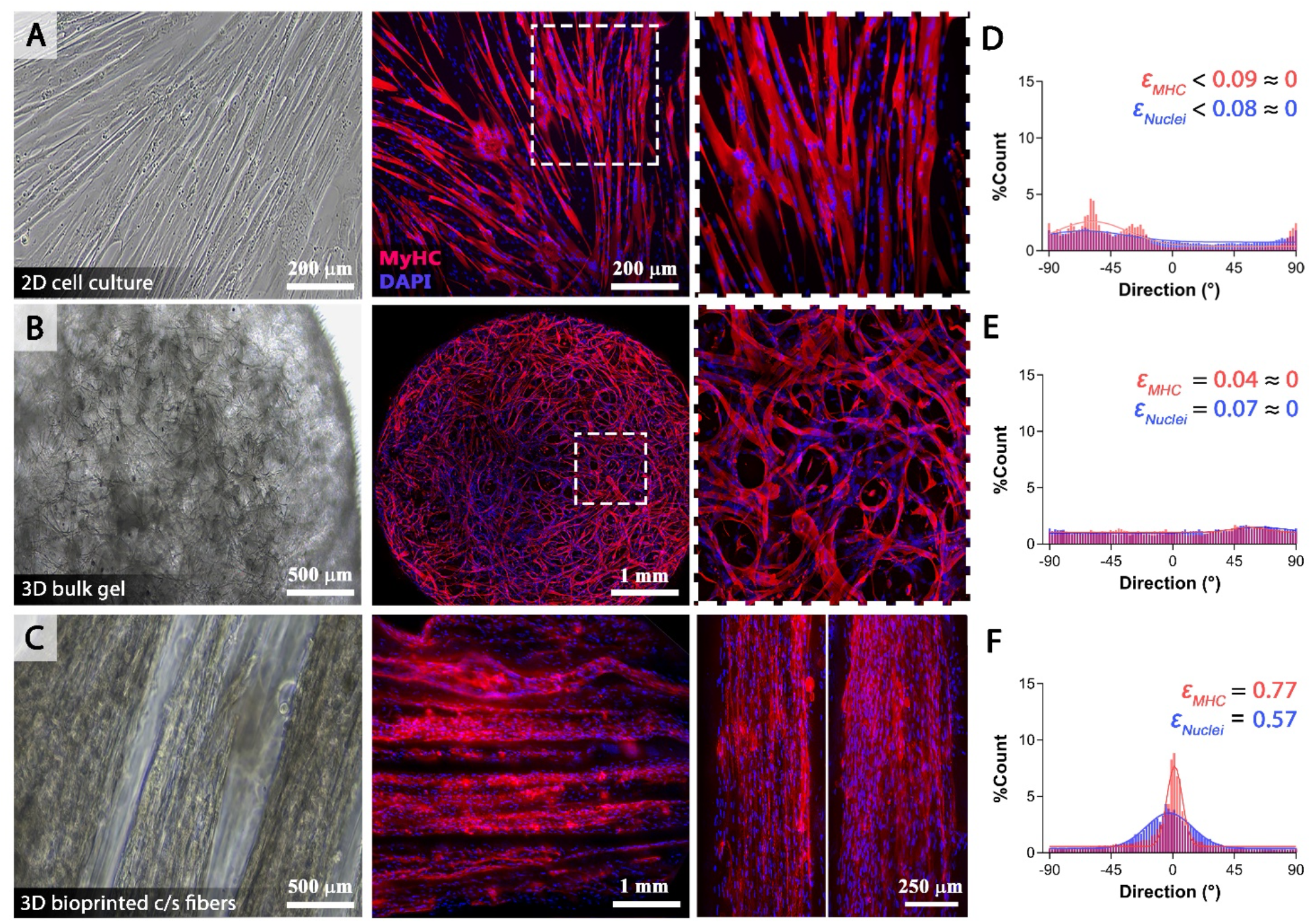
*In vitro* human pericyte (hPeri) myogenic structuration in the three studied groups: 2D, 3D bulk gel, and 3D bioprinted. A-C) Phase contrast and confocal images of hPeri cultured on a 2D substrate (A), in a 3D bulk gel (B) and in 3D bioprinted core/shell fibers (MyHC is shown in red while DAPI staining is shown in blue). D-F) Directionality analysis of the myo-substrates obtained in the three studied groups. Data are representative of at least three independent repeats.

The improved myotube alignment in the bioprinted group represents a positive, promising feedback, mimicking the cellular organization of native skeletal muscles. This feature can be appreciated from the directionality analysis performed on confocal fluorescence images (Figure 2D-F): while 2D and 3D bulk gel groups showed respectively localized or no alignment at all, bioprinted myo-structures revealed significantly sharper distributions centered along fiber major axis. To condense such differences into a practical value, data obtained from directionality analyses were fitted with a circular orientation distribution function (CODF, see Materials and Methods). Such function contains a single fit parameter 𝜀 that behaves in a similar way to an *order parameter*: a value close to 1 indicates strong alignment, while progressively smaller values indicate lesser alignment (for a random sample, the scattering is isotropic and 𝜀 = 0). As expected, 2D and 3D bulk groups returned an 𝜀 value close to 0, while biofabricated samples, on the other hand, were characterized by significant higher values (𝜀_MyHC_ = 0.77, 𝜀_nuclei_ = 0.57) confirming their high degree of alignment.

### 2.3 High-resolution mass spectrometry-based proteomic

As shown, skeletal muscle-derived human pericytes (hPeri) are a powerful source of competent myogenic precursor cells and such myogenic potential is favored by culturing these cells in 3D environments using PF matrix as scaffold. However, mechanistic information on how the PF environment rewires these cells to form functional muscle bundles is currently missing. To pinpoint molecular details at proteome resolution level, we decided to apply state-of-the-mass spectrometry-based proteomics to dissect molecular information from pericyte-based myo-substitutes. Specifically, we selected ALP+ hPeri from the skeletal muscle biopsies of four healthy donors to establish primary cultures of such progenitors. Next for each donor, we differentiated hPeri into mature myotubes by culturing these cells either in standard bi-dimensional (2D) cultures, in 3D bulk hydrogel or 3D-bioprinted aligned fibers via home-made core-shell 3D bioprinting (Figure 3A). After 20 days, hPeri gave rise to elongated MyHC positive syncytia (Figure 2) that were harvested and subjected to standard lysis procedures. In solution digestion coupled with a label-free liquid chromatography (LC)-MS/MS quantitation approach^16, 17^ allowed us to quantitate ∼3,200 proteins (Table S1). Proteome measurements were highly accurate and reproducible with a Pearson correlation coefficient among biological replicates ranging between 0.84 and 0.98 (Supplementary FigureS2A). Principal component analysis (PCA) revealed that the proteome profiles efficiently discriminate different samples according to their culture conditions (Figure 3B). The drivers of the separation between 2D cultures and 3D constructs (component 1 of the PCA loadings) were significantly enriched (FDR<0.05) for muscle- and extracellular matrix-related terms, confirming that 3D culture conditions favor differentiation and maturation of hPeri into contractile competent myofibers (Figure 3C).

**Figure 3.**
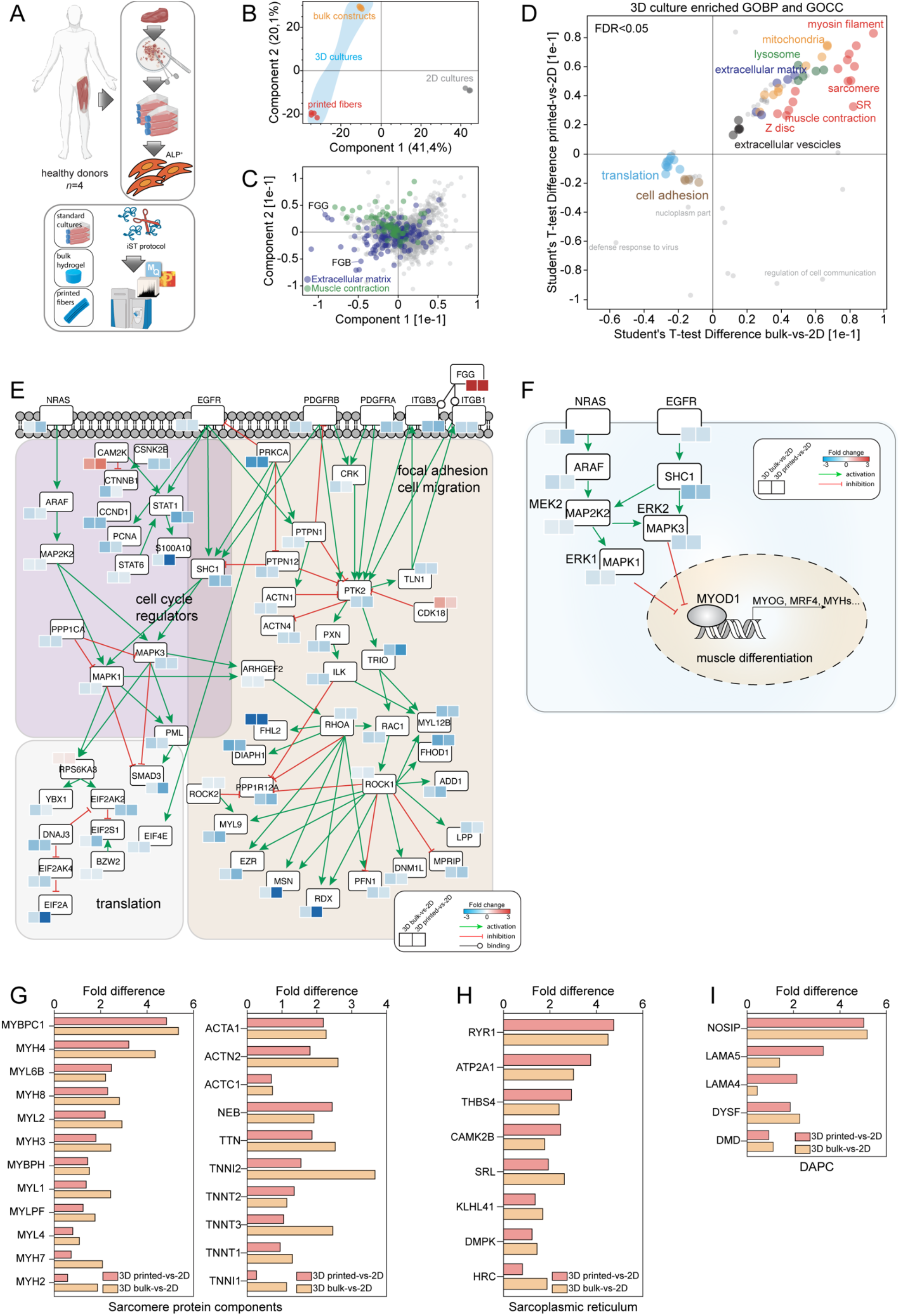
High-resolution proteomics unravels the proteomic asset of 2D- and 3D-cultured human pericytes. (A) Representative scheme summarizing the proteomic workflow. (B) Principal component analysis (PCA) showing sample grouping across the considered experimental conditions. (C) Scatter plot showing the major protein determinants driving sample segregation in the PCA plot. Proteins belonging to the extracellular matrix are highlighted in blue while those associated with muscle contraction are reported as green dots. (D) Two-dimensional annotation enrichment analysis of the significantly modulated proteins in Bioprinted/2D (y-axis) and bulk/2D (x-axis). Groups of related GO terms are labelled with the same color, as described in the inset. (E) Casual network summarizing the main molecular events occurring in human pericytes upon 3D differentiation. (F) Casual network reporting signaling relationship between mitogenic pathways and MYOD1. (G) Bar plot reporting the fold difference (3D-Bioprinted-VS-2D cultured and 3D-bulk-VS-2D cultured) of proteins annotated as “sarcomere protein components”. (H) Bar plot reporting the fold difference (3D-Bioprinted-VS-2D cultured and 3D-bulk-VS-2D cultured) of proteins annotated as “sarcoplasmic reticulum”. (I) Bar plot reporting the fold difference (3D-Bioprinted-VS-2D cultured and 3D-bulk-VS-2D cultured) of proteins annotated as “DAPC”.

**Figure 4.**
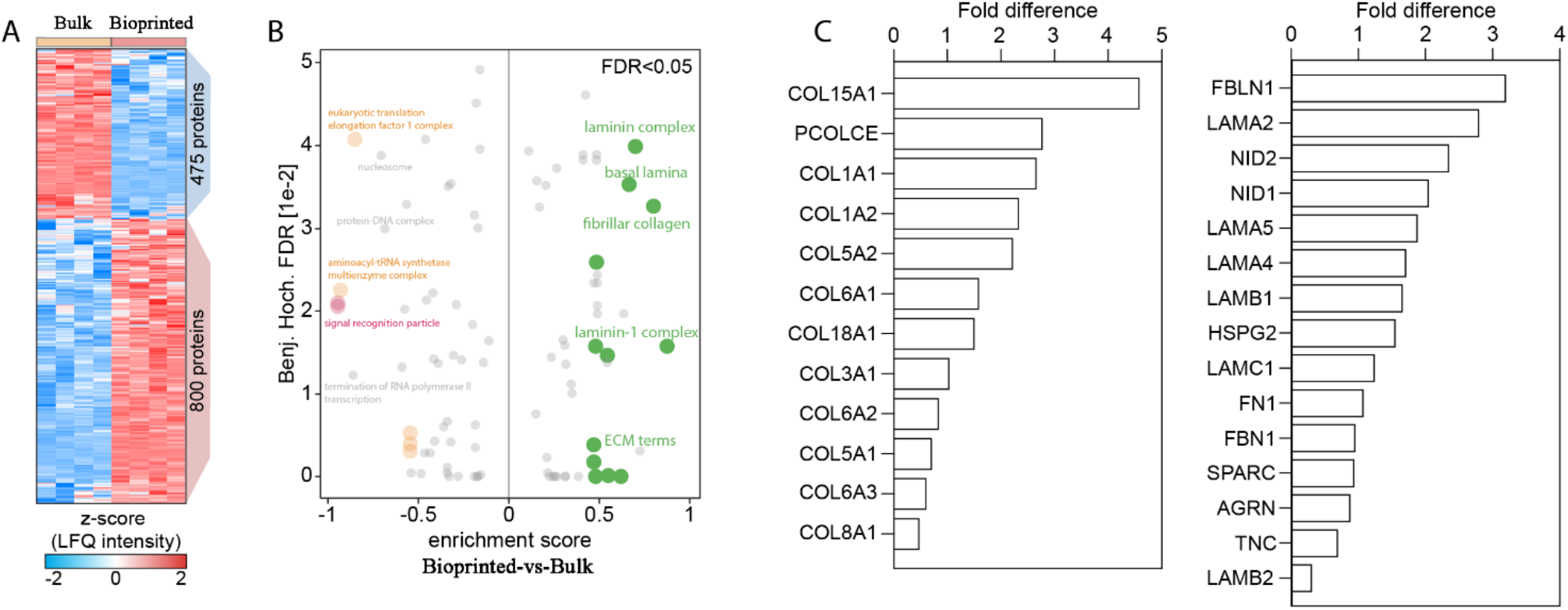
3D bioprinted hPeri are stimulated to create the muscle microenvironment. (A) Heatmap showing significantly modulated proteins in the 3D-Bioprinted-vs-3D-bulk comparisons. (B) One-dimensional annotation enrichment analysis of the significantly modulated proteins in 3D- Bioprinted-vs-3D-bulk comparisons. Similar terms are marked with the same color. (C) Bar plots reporting the fold difference in the 3D-Bioprinted-vs-3D-bulk comparisons of proteins annotated as “collagen” and “extracellular matrix constituent”.

PF biomimetic matrix profoundly changes the proteome of differentiated pericytes, approximately the ∼61% of the proteome of hPeri differentiated in bulk hydrogel constructs was found significantly different (FDR<0.01) when compared to cells differentiated in a 2D state, allowing the identification of 1956 differentially regulated proteins (471 up-regulated and 1485 down-regulated) (Supplementary FigureS2B). A slightly pronounced effect was found for hPeri differentiated in 3D-bioprinted fibers where proteome changes approaches ∼66% (FDR<0.01, 729 up-regulated and 1407 down-regulated) (Supplementary FigureS2B).

Pearson correlation (PC) coefficient between the proteome comparisons (Bulk-vs-2D and Bioprinted- vs-2D) approaches 0.7 (Supplementary FigureS2C) and includes 1577 significant proteins that are shared in both groups (Supplementary FigureS2D), suggesting that, independently from the degree of organization, PF hydrogels matrix may exert a conserved impact on pericyte cells. To investigate on this conserved effect, we used the two-dimensional annotation (2D) enrichment analysis ^18^ to classify processes and compartments that were significantly (FDR<0.05) represented/depleted in hPeri when differentiated in a PF hydrogel environment (Figure 3D). According to the gene ontology (GO) library, muscle-specific processes (red dots), mitochondria-(yellow dots), lysosome-(green dots), extracellular matrix-terms (purple dots) and extracellular vesicles were found overrepresented on hPeri differentiated (Figure 3D) in a PF-rich environment while process associated with cell adhesion and translation were among the most depleted terms (Figure 3D).

To rationally digest the content of information of our proteomic survey, we mapped the shared 1577 significantly modulated proteins onto a literature-derived networks of signaling and physical interactions extracted from the SIGNOR database with the aim to draw a PF-specific network. The resulting network encompasses 198 nodes including receptors and signal transducers while relationships between nodes are either positive (activations) or negative (inhibitions). Specifically, this strategy revealed that, in pericytes-derived myotubes, mitogenic pathways (RAS, EGFR, PGDFRA and PDGFRB) as well as focal adhesion components (ITGB1, ITGB3 and PTK2) and cytoskeleton remodeling protein effectors (ROCK1, RHOA and RAC1) were negatively affected in concentrations by the physical constrains created by the PF matrix (Figure 3E). To explain how such intracellular asset may contribute to stimulate pericyte myogenesis, we applied a transcription factor (TF) enrichment analysis to identify candidate myogenic downstream regulators that are sensitive of such low mitogenicity environment (Supplementary FigureS2E). We found that in both comparisons (Bulk-vs-2D and Bioprinted-vs-2D) the up-regulated proteins significantly enriched for all the members of the myogenic regulatory factors (MRFs), including MYOG, MYOD1, MYF6 and MYF5, that were among the top 10-ranked TFs (Supplementary FigureS2E). Mitogenic pathways including RAS-RAF-MAPK and EGFR signaling represses myogenic differentiation by inhibiting these TFs, especially MYOD and MYOG to ensure the efficient expansion of myoblasts pool ^19, 20^ (Figure 3F). Consistently, PF matrix derepresses such constrains by massively inducing the expression of all sarcomere proteins including myosin, actin and troponin isoforms (Figure 3G). Late myogenic genes, including sarcoplasmic reticulum proteins (Figure 3H) and components of the dystrophin associated glycoprotein complex were also increased in PF constructs (Figure 3I). Hence, for the very first time, our results demonstrated that PEG-Fibrinogen biomimetic matrix confers a low mitogenicity environment that favors, in an anchoring-independent three-dimensional state, the myogenic program stimulating the formation of contractile-competent bundles into pericytes-derived myotubes.

### 2.4 3D Bioprinted pericytes-derived myo-substitutes are stimulated to create the skeletal muscle niche

As shown in Figure 2, when extruded with the core/shell systems, pericyte cells are physically constrained to differentiate in highly aligned fibers, facilitating bundle orientation and myofiber maturation. By contrast, bulk constructs gave rise to randomly oriented mature myotubes. To uncover molecular differences coming from the two selected strategies for generating 3D human myosubstitutes, we compared the proteome profiles collected from the 3D bulk and 3D bioprinted samples with the aim to find protein signatures that may help to rationally describe, with molecular depth. Such approach identified 1275 significantly regulated proteins (800 up-regulated and 475 down-regulated proteins). We used the one-dimensional annotation (1-D) enrichment analysis to identify significantly (FDR<0.05) enriched processes in the considered comparison.

According to the gene ontology (GO) library, basal lamina- and ECM-related terms (green dots) were the most enriched terms that describe differences among 3D bulk and 3D bioprinted constructs. Consistently, collagen and fibrillar components are largely induced in biofabricated PF fibers. This suggests that in this physically constrained environment pericytes are stimulated to produced basal lamina components that act as a scaffold for pericyte myogenesis, thus recreating the physiological skeletal muscle fiber niche. Thus, besides the effect of 3D PF environment increasing the differentiation efficacy of hPeri by promoting the formation of contractile-competent bundles, biofabrication strategy remarkably affect hPeri myogenic outcome stimulating the production of ECM specific components (Laminin, Basal Lamina, Laminin-1 complex, etc.) fundamental for establishing the physiological architecture of skeletal muscle tissue. As a result, the suggested 3D rotary wet-spinning (ROWS) biofabrication method proves to be a really effective strategy to manufacture structurally and functionally equivalent human-derived myo-substitutes.

### 2.5 hPeri-derived 3D bioprinted myo-substitute implantation into a VML mouse model

We evaluated the potential of our biofabrication strategy by grafting a human pericyte-derived myo-substitute into a volumetric muscle loss (VML) animal model. Prior to transplantation, we ensured that the 3D bioprinted myo-substitute exhibited essential characteristics, including the production of matrix, sarcomeric proteins, and muscle-specific proteins. Our VML mouse model consisted in the surgical removal of 30% of the Anterior Tibialis (TA) muscle, following a *tried-and-true* protocol ^2, 14^ (see Figure 5). The chosen degree of muscle ablation (30%) aimed to avoid xenograft issues, as even immunocompromised mouse strains still possess active macrophages that may rapidly invade the ablated TA and target *non-self* cells. Moreover, the selection of the damage size for VML modeling was based on previously published data, which reported the critical size of muscle defect in order to generate a reliable VML damage model ^2, 14, 21, 22^. The hPeri-derived myo-substitutes were grafted into the site after being appropriately trimmed to fit the TA lodge created by the removed muscular tissue. The contralateral TA served as the sham control. After a 30-day period post-implantation, the sham control group showed no signs of regenerating or replenishing tissue in the damaged area, as depicted in Figure 5B, C. In contrast, the TAs that had been treated with the RoWS biofabricated structure exhibited evidence of hPeri-derived regenerating tissue, as observed in Figure 5D, E. The center nucleated muscle fibers within the regenerated tissue displayed positive staining for the human-specific markers LaminA/C and Myosin Heavy Chain (as indicated by the arrows in Figure 5F, G). These results highlight the ability of the hPeri-bioprinted myo-substitute to produce muscle tissue capable of replacing the removed TA mass and closing the gap created by the VML. Furthermore, we assessed the vascularization potential of the RoWS-generated myo-structures by performing immunofluorescence staining against the vessel-specific marker Smooth Muscle Actin (SMA), which labels the blood vessel wall. The immunofluorescence images, shown in Figure 5F, Ii, provided insights into the engraftment capability of the RoWS-generated structures, confirming the presence of a neo-vascularization process within the repaired tissue.

**Figure 5.**
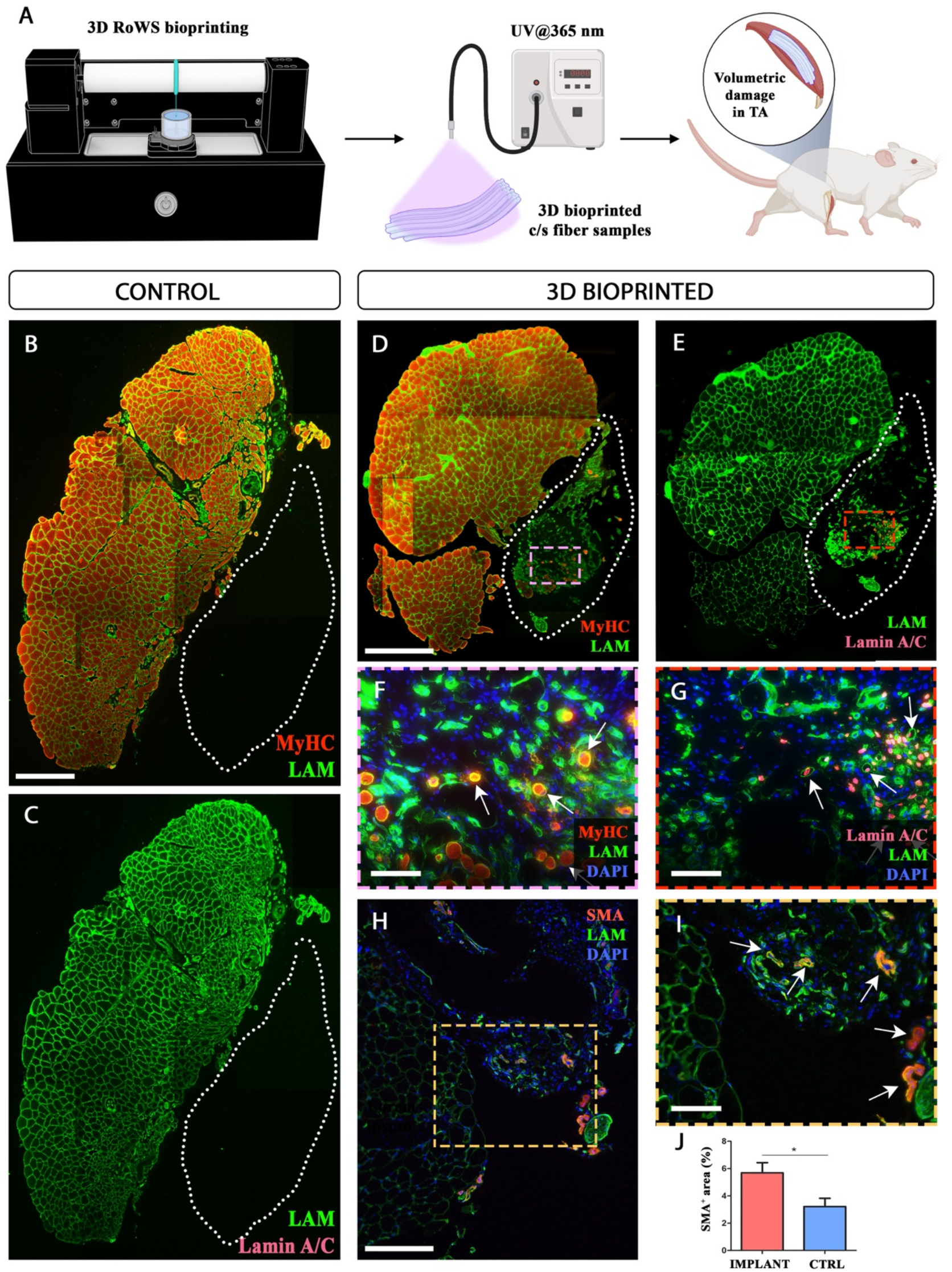
hPeri derived 3D bioprinted myo-substitute engraftment into a mouse TA volumetric damage. a) A) Schematic representation of the 3D RoWS derived myo-substitute implantation. Reconstructed TA cross-sections from B, C) sham (control) and D-I implanted (treated with 3D bioprinted myosubstitutes) mice; immunofluorescence against Myosin Heavy Chain (MyHC), Laminin (LAM) and human Lamin A/C (Lamin A/C). F, G) Enlarged views of selected areas respectively from (D) and (E). H) Smooth Muscle Actin (SMA) immunostaining at the interface between implant and host native muscle demonstrating the neo-vascularization process occurring in the implanted mice. I) Magnification of a selected area from (H) showing the 3D RoWS derived myo- substitute integration in the host muscle tissue. J) Quantification of SMA positive areas in the cross-sections obtained from the implant and control samples. Scale bars: B=150μm; D=200μm, F, G, I=50 μm; H=100μm.

The regenerating area showed a remarkable recruitment of SMA positive blood vessel, pivotal for the engraftment and survival of the implanted myo-substitute. In the enlarged view of the grafted area, it is appreciable the infiltration of small and larger vessel (Figure 5I). Hence, exploiting image J software function it has been carried out vascularization quantification demonstrating a remarkable vessel colonization in the pericyte derived reconstructed muscle tissue, displaying twice of the SMA positive area comparing with the TA undamaged area as control (Figure 5J). Since, 3D bioprinted hPeri myo-substitute demonstrated an important engraftment capacity recruiting blood vessel, integrating into host TA muscle and generating human derived myofibers.

It is well-known pericytes ability to secrete a wide range of soluble mediators that contribute to vessel stability and recruitment in angiogenesis processes ^23, 24^. Considering the potential influence of the 3D environment on pericyte behavior, we investigated our proteomics survey to identify candidate angiogenic molecules that exhibit significantly increased abundance in the 3D groups. To accomplish this, we constructed a protein ranking system that prioritized significant proteins based on their differences in abundance between the 3D bulk-vs-2D and 3D bioprinted-vs-2D comparisons. We specifically focused on proteins annotated with the term “angiogenesis” in the Gene Ontology Biological Process (GOBP) library (see Supplementary FigureS3A).

Among the top-ranked angiogenic factors identified, Thrombospondin-4 (THBS4) stood out as the only macromolecule with recognized pro-angiogenic properties that exhibited conservation in both group comparisons (Supplementary FigureS3A). The human genome contains five THBS genes that encode secreted glycoproteins, whose expression is induced in sites of tissue damage or active remodeling ^25^. THBS4, in particular, plays a crucial role in regulating physical interactions between cells and the extracellular matrix, making it a potent angiogenic factor^25, 26^. This study represents the first identification of a molecular candidate responsible for the pericyte’s angiogenic capabilities, specifically in the context of vessel recruitment during muscle reconstruction under the influence of a 3D biofabricated environment.

## 3. Conclusions

Skeletal muscle (SM) tissue is a peculiar example of how intertwined the structural and functional properties of a tissue can be, from the molecular level up to the macro scale. Regrettably, such orchestrated organization is loss in several debilitating conditions including traumatic injuries, invasive surgeries or devastating muscle diseases. In those cases, the massive removal/wasting of functional muscles (i.e. volumetric muscle loss or VML) is not compensated by the intrinsic regenerative potential of this tissue. Fortunately, current advances in biotechnology and bioengineering are paving the way for a new “regenerative” era. Novel platforms and bioproducts (i.e. bioreactors, bioinks and bioprinting systems) are opening at unprecedent level the possibility to generate miniaturized, and in most cases at normal scale, functional organs that can be used to treat or to replace the damaged ones. In the case of muscle biology, the major challenge is to develop and validate a system capable to recapitulate its complex architecture composed by hierarchical compartment of connective tissue and muscle cells, while demonstrating the full biocompatibility and grafting of such constructs in host recipients.

To this end, in the present manuscript, we proposed a biofabrication strategy that leverages wet-spinning principles to create anisotropic 3D constructs comprising densely packed, parallelly aligned hydrogel fibers. Through the incorporation of a custom microfluidic printing head, we achieved continuous fabrication of core/shell fibers. By optimizing the chemical composition and mechanical properties of these fibers, we successfully induced the robust differentiation of human-derived skeletal muscle progenitors, leading to the formation of well-organized myotubes and the expression of important marker proteins like MHC.

While, for the very first time, here we demonstrated by means of state-of-the-art mass spectrometry-based proteomics the effect of 3D PEG-Fibrinogen matrix providing a low mitogenicity environment besides blocking cell adhesion machinery. Thus, promoting the myogenic program by facilitating the formation of contractile-competent bundles in pericyte-derived myotubes. Following *in vitro* investigation, we tested our biofabricated myo-substitutes within a VML animal model. The biofabricated structures revealed the ability to regenerate muscle tissue, as evidenced by the presence of hPeri-derived regenerating tissue containing positive markers for human-specific proteins, and overall to integrate into host muscular tissue. Vascularization assessment further demonstrated optimal grafting e biocompatibility of RoWS-generated structures with the host recipients. While the main limitation of this work is the absence of functional assessments of RoWS-generated structures, the superior contraction ability of tightly packed myobundles infers the almost complete sarcomere functions of these 3D-bioprinted myo-substitutes. Of note, by investigating our proteomic survey, we identified THBS4 as a key angiogenic factor that exhibits upregulation in the 3D environment created through the proposed RoWS biofabrication process. We speculate that the upregulation of THBS4 may contribute significantly to the successful engraftment of the myo-substitute into the host muscle. Future studies should further investigate this aspect to elucidate this intricate process.

## 4. Materials and Methods

### Materials

Unless otherwise noted, all chemicals were obtained from Sigma-Aldrich and used directly. FMC Biopolymers generously provided sodium alginates with low molecular weights (LMW-ALG, Mw 33 kDa) and high molecular weights (HMW-ALG, Mw 100 kDa). ALG-FITC and ALG-TRITC were synthesized following a previously published procedure^15^.

### 3D wet-spinning bioprinting setup

Using a customized 3D rotary wet-spinning bioprinter developed in our lab, core/shell hydrogel fibers were created and spatially deposited in three dimensions. The entire platform includes an extrusion system made up of a rotating drum collector with a diameter of 20 mm and a length of 180 mm, an X-axis with a travel range of 160 mm, and a crosslinking bath microtank with a coaxial nozzle placed at the bottom for the immediate gelation of extruded core/shell fibers (Inner needle diameter: Outer needle diameter: 500 m). Using an Arduino Mega board and specialized Python software, the entire system was controlled. Utilizing a transparent and class I biocompatible resin (Class I polymer for surgical guides as per Rule 5, Annex IX of Medical Devices Directive 93/42/EEC) and stereolithography 3D printing (DWS 3500PD), the microfluidic printing head was created. The nozzle was thoroughly rinsed in isopropanol to remove unreacted photoresin that was trapped in the channels after the 3D printing, dried, and then crosslinked for an additional 10 minutes in a UV curing oven. The 3D printed MPH was then mounted on the x-axis arm of the bioprinter and given a crosslinking bath microtank.

### Human Pericytes (hPeri) isolation procedure and culture

hPeri were isolated from muscle biopsies collected from healthy donors. Tissues were minced and then digested with 100U/ml of Collagenase Type II (Gibco, Life Technologies, Carlsbad, CA, USA) in Phosphate Buffered Saline (PBS) with Ca^2^ and Mg^2^, for 60 min. Afterward, muscle fragments were gently dissociated by pipetting and passing through a 100μm cell strainer. Cell suspension was centrifuged for 10’ at 1200 rpm, and the pellet was resuspended and filtered again with a 70μm cell strainer. Cell suspension was then centrifuged for 10’ at 1200 rpm, and the pellet was resuspended and seeded 100-mm Petri dishes at low confluences (1.000 cells/cm2) in CYTO-GROW medium (Resnova), supplemented with 100 IU/mL penicillin and 100 mg/mL streptomycin (Gibco). Cells were expanded and cultured in 100-mm Petri dishes.

### *In vitro* analysis of hPeri myogenic structuration in 2D and 3D culture conditions

To study hPeri myogenic structuration, the cells were plated in 6 well Petri dishes (3×10^5^) or used in bioink formulation with the aim to produce 2D cell culture, 3D bulk and 3D RoWS derived myo**-** constructs. hPeri derived cultures were grown up to 21 days in CYTO-GROW (Resnova) supplemented with 100 U/mL penicillin and 100 mg/mL streptomycin (Gibco) at 37 °C and 5% CO2 humid atmosphere until reaching the complete differentiation condition.

### Fabrication of human pericytes-loaded 3D myogenic structures

#### 3D Bulk structures

3D bulk hydrogels were fabricated by a casting method: human derived pericytes 2 x 10^7^ hPeri/mL were soaked into a sterile-filtered precursor solution made of 0.8% w/v PEG-Fibrinogen in 25 mM HEPES buffer and 0.1% w/v Irgacure 2959 as radical photoinitiator. Afterwards, the mixture was poured into cylindrical silicon molds (PDMS) and then exposed for 5 minutes to UV light for PEG-Fibrinogen polymerizing, the bulk structures were placed in CYTO-GROW (Resnova) supplemented with 100 U/mL penicillin and 100 mg/mL streptomycin (Gibco) at 37 °C and 5% CO2 humid atmosphere.

#### 3D bioprinted core/shell fiber structures

The custom co-axial MPH was used to create core/shell hydrogel fibres. The core bioink contained 2 x 10^7^ hPeri/mL suspended in a sterile-filtered solution containing 0.2% w/v LMW-ALG and 0.8% w/v PEG-Fibrinogen in 25 mM HEPES buffer solution. In order to create the shell biomaterial ink, 2% w/v LMW-ALG and 1% w/v HMW-ALG were dissolved in 25 mM HEPES. The MPH and tubing were cleaned with a 70% ethanol solution prior to use, and then flushed with sterile dH2O. To create the co-flow in the extrusion nozzle, the inks were fed continuously into the MPH nozzle. The typical core and shell flow rates in all cellular experiments were Q_c_ = 160 mL/min and Q_s_ = 320 mL/min. After that, 0.6 M CaCl2 solution was added to the crosslinking bath microtank before the extrusion of the core and shell inks. Upon coming into contact with CaCl2, the core/shell fibres immediately begin to gel in the vicinity of the nozzle’s tip. A tweezer is used to gently pull the hydrogel fibre that results up until it touches the Teflon drum’s surface. A hydrogel fibre begins to continuously extrude from the nozzle and collect onto the drum in the form of a bundle as soon as the fibre contacts the drum. The drum’s rotational speed was set to 64 rpm, and the number of threads in each bundle (37 threads per bundle, 30 seconds of extrusion) remained constant. The samples were taken from the rotating drum after extrusion. The bundles were exposed to 365nm UV light for 5 minutes in order to crosslink the bioink in the core. Following this, the bundles were placed in CYTO-GROW (Resnova) supplemented with 100 U/mL penicillin and 100 mg/mL streptomycin (Gibco) at 37 °C and 5% CO2 humid atmosphere.

### Skeletal muscle organization image analysis

Thanks to its unique aligned architecture, we estimated the organization of SM artificial tissue in the studied groups by computing the directionality of immunofluorescence images using Fiji built-in plugin (https://imagej.net/plugins/directionality). Data obtained from directionality analyses were then fitted with a circular orientation distribution function (CODF, 𝐼(𝜃) = (1 - 𝜀²)/[(1 + 𝜀)² - 4𝜀 cos² 𝜃]) where the single fit parameter 𝜀 behaves in a similar way to an order parameter: a value close to 1 indicates strong alignment, while progressively smaller values indicate lesser alignment. For a random sample, the scattering is isotropic and 𝜀 = 0. Interestingly, both DAPI and MHC channels were used in the analysis, providing insights about the alignment of nuclei and myotubes.

### Proteome sample preparation

Samples were harvested and directly lysed in ice-cold RIPA buffer. In-gel fractionation was performed via SDS-PAGE electrophoresis. Gel lanes were cut into equal pieces and digested. In brief, gel pieces were washed, de-stained and dehydrated. Proteins were reduced with 10 mM dithiothreitol (DTT), alkylated with 55 mM iodoacetamide (IAA) and digested with the endopeptidase sequencing-grade Trypsin (Promega; Madison, WI, USA) overnight at 37°C. Collected peptide mixtures were concentrated and desalted using via in-StageTip (iST) method^27–29^. Samples were separated by high-performance liquid chromatography in a single run (without pre-fractionations) and analyzed by LC-MS/MS.

### LC-MS/MS measurements

Instruments for LC-MS/MS analysis consisted of a NanoLC 1200 (Thermo Fisher Scientific) coupled via a nano-electrospray ionization source to a quadrupole-based Q Exactive HF benchtop mass spectrometer (Thermo Fisher Scientific). Peptide separation was carried out according to hydrophobicity on a home-made column [75 μm inner diameter, 8 μm tip, 400 mm bed packed with Reprosil-PUR, C18-AQ, 1.9 μm particle size, 120 Å pore size (New Objective, PF7508-250H363)], using a binary buffer system consisting of solution A (0.1% formic acid) and B (80% acetonitrile, 0.1% formic acid). Total flow rate was 300nl/min. For the liquid chromatography linear gradient, after sample loading, the run started at 5% buffer B for 5 min, followed by a series of linear gradients, from 5% to 30% B in 90 min, then a 10 min step to reach 50% and a 5 min step to reach 95%. This last step was maintained for 10 min.

MS spectra were acquired using 3×10^6^ as an automatic gain control (AGC) target, a maximal injection time of 20 ms and 120,000 resolution at 200 m/z. The mass spectrometer operated in data-dependent Top20 mode with subsequent acquisition of higher-energy collisional dissociation fragmentation MS/MS spectra of the top 20 most intense peaks. Resolution for MS/MS spectra was set to 15,000 at 200 m/z, AGC target to 1×10^5^, maximum injection time to 20 ms and the isolation window to 1.6 Th. The intensity threshold was set at 2.0×10^4^ and dynamic exclusion at 30 s.

### Proteome data processing

All acquired raw files were processed using MaxQuant (1.6.2.10) and the implemented Andromeda search engine. For protein assignment, spectra were correlated with the Human (v. 2021) reference proteome, including a list of common contaminants. Searches were performed with tryptic specifications and default settings for mass tolerances for MS and MS/MS spectra. The other parameters were set as follows: fixed modification, carbamidomethyl (C); variable modifications, oxidation, acetyl (N-term); digestion, trypsin, Lys-C; minimum peptide length, 7; maximum peptide mass, 470 Da; false discovery rate for proteins and peptide spectrum, 1%.

For further analysis, Perseus software (1.6.2.3)^18^ was used, and first filtered for contaminants and reverse entries as well as proteins that were only identified by a modified peptide (first filter). The label-free quantitation (LFQ) ratios were logarithmized, grouped and filtered for minimum valid number (minimum of three in at least one group) (second filter).

Missing values were replaced by random numbers drawn from a normal distribution. Two-sample t-test analysis was performed using S0=0.1 FDR<0.01. Proteins with Log2 difference ≥±1 and q-value <0.01 were considered significantly enriched. Categorical annotation was added in Perseus in the form of GO biological process, GO molecular function and GO cellular component, and Kyoto Encyclopedia of Genes and Genomes (KEGG) pathways.

### Protein network generation

This strategy has been previously developed and applied to query complex muscle-specific proteome datasets^29, 30^. Casual relationships between significant protein entities were retrieved from the SIGNOR database^31^ using a dedicated application in the Cytoscape platform^32^. Nodes were color coded according to their difference value in the dedicated comparison.

### *In vivo* 3D RoWS derived myo-substitute implantation in a VML mouse model

Two-month-old male SCID/Beige mice were anesthetized with an intramuscular injection of physiologic saline (10 ml/kg) containing ketamine (5 mg/ml) and xylazine (1 mg/ml) and then the 3D 3D RoWS derived myo**-**constructs were implanted in tibialis anterior muscle (TA) of a VML mouse model recreated according to following surgical procedure ^2, 14^: (i) To get as far as the TA a confined incision on the medial side of the leg has been executed (ii) the muscle fibers were deeply removed (leaving in place the tendons) by employing a cautery to avoid bleeding and to create an adequate shelter for the implant, and (iii) the bioprinted myo-substitute was positioned in the removed TA fibers lodge and the incision was sutured. In the contralateral TA, used as control, no construct was implanted, it was only surgically ablated and sutured to produce a VML damage.

Analgesic treatment (Rimadyl, Pfizer, USA) was administered after the surgery to reduce pain and discomfort. Mice were sacrificed 30 days after implantation for molecular and morphological analysis. Experiments on animals were conducted according to the rules of good animal experimentation I.A.C.U.C. No 432 of March 12, 2006, and under Italian Health Ministry approval n° 228/2015-PR.

### Immunofluorescence

Following in vitro culture, 2D 3D bulk and 3D RoWS derived myo-constructs were fixed in 2% PFA and processed for fluorescence microscopy as previously described ^14^.

The in vitro constructs were incubated with anti-Myosin Heavy Chain (MHC, Clone MF20 DSHB, 1:50 titer). Samples were counterstained with 4′,6-diamidino-2-phenylindole (DAPI) to detect nuclei and mounted on glass slides with glycerol to be imaged with Nikon A1R laser scanning confocal microscope with the NIS software.

Conversely, constructs explanted from SCID mice were embedded in O.C.T. and quickly frozen in liquid nitrogen cooled isopentane for sectioning at a thickness of 8 mm on a EPREDIA cryostat. Sections were permeabilized with 0.3% Triton X-100 in PBS for 30 min and blocked in blocking solution formed by 10% of goat serum, 1% of glycine, 0,1% triton X-100 in PBS for 1h at RT. Subsequently, sections were incubated with primary antibody in blocking solution for 2h at RT. Primary antibody were diluted as follows: mouse monoclonal anti-Myosin Heavy Chain (MHC, Clone MF20 DSHB) 1:2, rabbit polyclonal anti-Laminin (LAM, Sigma-Aldrich) 1:200, mouse monoclonal anti-a-smooth muscle actin (SMA, Sigma-Aldrich) 1:100, rabbit polyclonal anti von Willebrand factor (vWF, Abcam) 1:100, mouse monoclonal anti-Lamin A/C (Thermo Fisher Scientific) 1:200. After several washes with washing solution formed by 1% BSA and 0,2% Triton X-100 in PBS, sections were incubated with Alexa Fluor 555-coniugated goat anti-mouse IgG (H+L; Thermo Fisher Scientific, 1:400) and Alexa Fluor 488-coniugated goat anti-rabbit (H+L; Thermo Fisher Scientific, 1:400) in blocking solution for 1 h at RT. Samples were counterstained with DAPI to detect nuclei, washed 3 times with PBS, and mounted on glass slides with Vectashield mounting medium (Vector Laboratories). Samples were imaged with a Nikon Eclipse TE2000 microscope equipped with a CoolSNAP MYO CCD camera (Photometrix) and MetaMorph software.

### Data availability

The mass spectrometry proteomics data are available at the ProteomeXchange Consortium via the PRIDE partner repository with the dataset identifier PXD043605.

## Funding

This work was supported by AFM-Téléthon (23551 to A.R.), Fondazione Telethon (TMPGMFU22TT to P.G.), Muscular Dystrophy Association (MDA 968551 to P.G.), National Science Centre Poland (NCN) within SONATA 14 (2018/31/D/ST8/03647 to MC and N.C.) and Ministero dell’Istruzione, dell’Università e della Ricerca (PRIN Funding Scheme No. 201742SBXA_004 to C.G.).

## Notes

### Competing Interest Statement

The authors have declared no competing interest.

## Bibliography

1. Kim, H., Osaki, T., Kamm, R. D. & Asada, H. H. Multiscale engineered human skeletal muscles with perfusable vasculature and microvascular network recapitulating the fluid compartments. Biofabrication 15, 15005 (2022).

2. Costantini, M., et al. Biofabricating murine and human myo-substitutes for rapid volumetric muscle loss restoration. EMBO Mol. Med. 13, e12778 (2021).

3. Fornetti, E. et al. A novel extrusion-based 3D bioprinting system for skeletal muscle tissue engineering. Biofabrication 15, 25009 (2023).

4. Volpi, M., Paradiso, A., Costantini, M. & Świȩszkowski, W. Hydrogel-Based Fiber Biofabrication Techniques for Skeletal Muscle Tissue Engineering. ACS Biomater. Sci. Eng. (2022).

5. Fornetti, E. et al. Dystrophic Muscle Affects Motoneuron Axon Outgrowth and NMJ Assembly. *Adv*. Mater. Technol. 7, 2101216 (2022).

6. Leng, Y. et al. Advances in In vitro Models of Neuromuscular Junction: Focusing on Organ-on-a-chip, Organoids and Biohybrid Robotics. Adv. Mater. 2211059 (2023).

7. Ladd, M. R., Lee, S. J., Stitzel, J. D., Atala, A. & Yoo, J. J. Co-electrospun dual scaffolding system with potential for muscle–tendon junction tissue engineering. Biomaterials 32, 1549– 1559 (2011).

8. Bakooshli, M. A., et al. A three-dimensional culture model of innervated human skeletal muscle enables studies of the adult neuromuscular junction and disease modeling. BioRxiv 275545 (2018).

9. Jeong, D. et al. Efficient Myogenic/Adipogenic Transdifferentiation of Bovine Fibroblasts in a 3D Bioprinting System for Steak-Type Cultured Meat Production. Adv. Sci. 9, 2202877 (2022).

10. Shahin-Shamsabadi, A. & Selvaganapathy, P. R. Engineering murine adipocytes and skeletal muscle cells in meat-like constructs using self-assembled layer-by-layer biofabrication: A platform for development of cultivated meat. Cells Tissues Organs 211, 304–312 (2022).

11. Dellavalle, A. et al. Pericytes of human skeletal muscle are myogenic precursors distinct from satellite cells. Nat. Cell Biol. 9, 255–267 (2007).

12. Fuoco, C., et al. Injectable polyethylene glycol-fibrinogen hydrogel adjuvant improves survival and differentiation of transplanted mesoangioblasts in acute and chronic skeletal-muscle degeneration. Skelet. Muscle 2, (2012).

13. Zhu, P. et al. Selective expansion of skeletal muscle stem cells from bulk muscle cells in soft three-dimensional fibrin gel. Stem Cells Transl. Med. 6, 1412–1423 (2017).

14. Fuoco, C. et al. In vivo generation of a mature and functional artificial skeletal muscle. EMBO Mol. Med. 7, 411–422 (2015).

15. Costantini, M. et al. Microfluidic-enhanced 3D bioprinting of aligned myoblast-laden hydrogels leads to functionally organized myofibers in vitro and in vivo. Biomaterials 131, 98–110 (2017).

16. Reggio, A. et al. Metabolic reprogramming of fibro/adipogenic progenitors facilitates muscle regeneration. Life Sci. Alliance 3, e202000646 (2020).

17. Reggio, A. et al. Development of a platform of 3D adipogenesis to model, at higher scale, the impact of LY2090314 compound on fibro/adipogenic progenitor adipogenic drift. Dis. Model. Mech. (2023) doi:10.1242/dmm.049915.

18. Tyanova, S. et al. The Perseus computational platform for comprehensive analysis of (prote)omics data. Nat. Methods 13, 731–740 (2016).

19. Dorman, C. M. & Johnson, S. E. Activated Raf inhibits avian myogenesis through a MAPK-dependent mechanism. Oncogene 18, 5167–5176 (1999).

20. Tortorella, L. L., Milasincic, D. J. & Pilch, P. F. Critical proliferation-independent window for basic fibroblast growth factor repression of myogenesis via the p42/p44 MAPK signaling pathway. J. Biol. Chem. 276, 13709–13717 (2001).

21. Turner, N. J. & Badylak, S. F. Regeneration of skeletal muscle. Cell Tissue Res. 347, 759– 774 (2012).

22. Anderson, S. E. et al. Determination of a critical size threshold for volumetric muscle loss in the mouse quadriceps. Tissue Eng. Part C Methods 25, 59–70 (2019).

23. Avolio, E. et al. Combined intramyocardial delivery of human pericytes and cardiac stem cells additively improves the healing of mouse infarcted hearts through stimulation of vascular and muscular repair. Circ. Res. 116, e81–e94 (2015).

24. Vono, R. et al. Activation of the pro-oxidant PKCβII-p66Shc signaling pathway contributes to pericyte dysfunction in skeletal muscles of patients with diabetes with critical limb ischemia. Diabetes 65, 3691–3704 (2016).

25. Peisker, F. et al. Mapping the cardiac vascular niche in heart failure. Nat. Commun. 13, 3027 (2022).

26. Muppala, S. et al. Thrombospondin-4 mediates TGF-β-induced angiogenesis. Oncogene 36, 5189–5198 (2017).

27. Kulak, N. A., Pichler, G., Paron, I., Nagaraj, N. & Mann, M. Minimal, encapsulated proteomic-sample processing applied to copy-number estimation in eukaryotic cells. Nat. Methods 11, 319–324 (2014).

28. Reggio, A. et al. Role of FAM134 paralogues in endoplasmic reticulum remodeling, ER-phagy, and Collagen quality control. EMBO Rep. 1–20 (2021) doi:10.15252/embr.202052289.

29. Reggio, A. et al. A 3D adipogenesis platform to study the fate of fibro/adipogenic progenitors in muscular dystrophies. Dis. Model. Mech. 16, (2023).

30. Reggio, A. et al. Metabolic Reprogramming of Fibro/Adipogenic Progenitors Facilitates Muscle Regeneration. SSRN Electron. J. (2019) doi:10.2139/ssrn.3411252.

31. Lo Surdo, P., et al. SIGNOR 3.0, the SIGnaling network open resource 3.0: 2022 update. Nucleic Acids Res. (2022) doi:10.1093/nar/gkac883.

32. Shannon, P. et al. Cytoscape: a software environment for integrated models of biomolecular interaction networks. Genome Res. 13, 2498–504 (2003).

