## Supplementary information for "3D bioprinting directly affects proteomic signature and myogenic maturation in muscle pericytes-derived human myo-substitute"

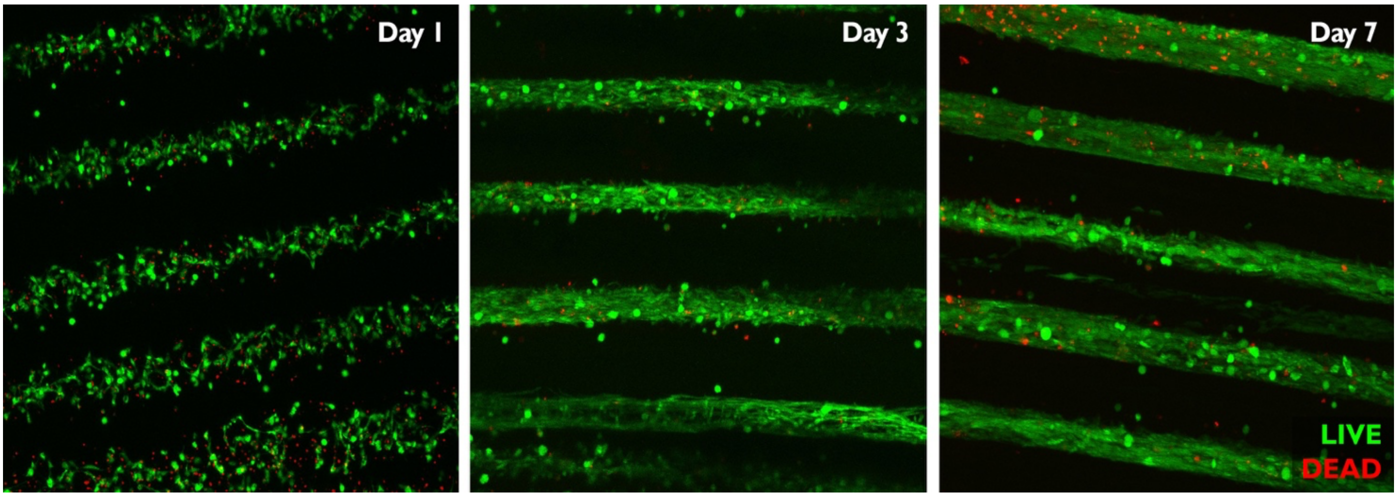


**Supplementary FigureS1**. **Live/Dead assay at various time points of pericyte-loaded 3D bioprinted core/shell fibers.** Live/dead staining of hPeri extruded with the core-shell bioprining system, cells were monitored at 1,3, and 7 days upon printing. The great majority of the cells was alive for all the studied time points; moreover, starting at day 3, one can appreciate the alignment of cells within the soft fiber cores in the extrusion direction. At day 7, the presence of viable myotubes can also be noticed. Images are representative of three independent biological repeats.

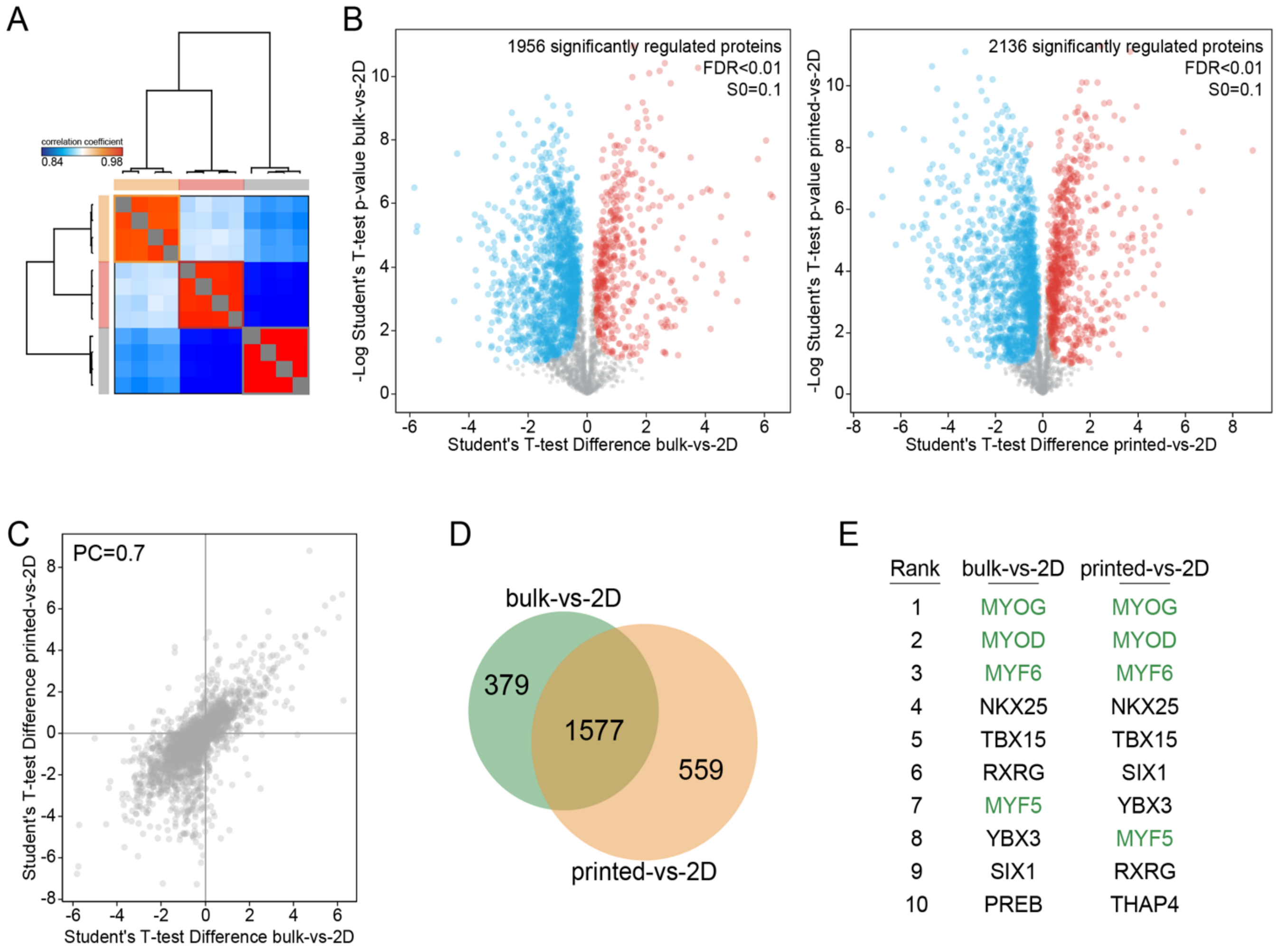


**Supplementary FigureS2**. **Proteomics of human pericytes**. (A) Heat map showing sample correlation between biological repeats. (B) Volcano plots reporting significantly regulated proteins (FDR<0.01) in the 3D-bulk-VS-2D (left) and the 3D-printed-VS-2D (right) comparisons. (C) Scatter plot showing protein correlation (PC=0.7) between significantly regulated protein coming from the 3D-bulk-VS-2D and the 3D-printed-VS-2D comparisons. (D) Veen diagram showing exclusive and commonly shared proteins in the 3D-bulk-VS-2D and the 3D-printed-VS-2D comparisons. (E) Transcription factor (TF) enrichment analysis reporting the relative TF ranking in the 3D-bulk-VS-2D and the 3D-printed-VS-2D comparisons.

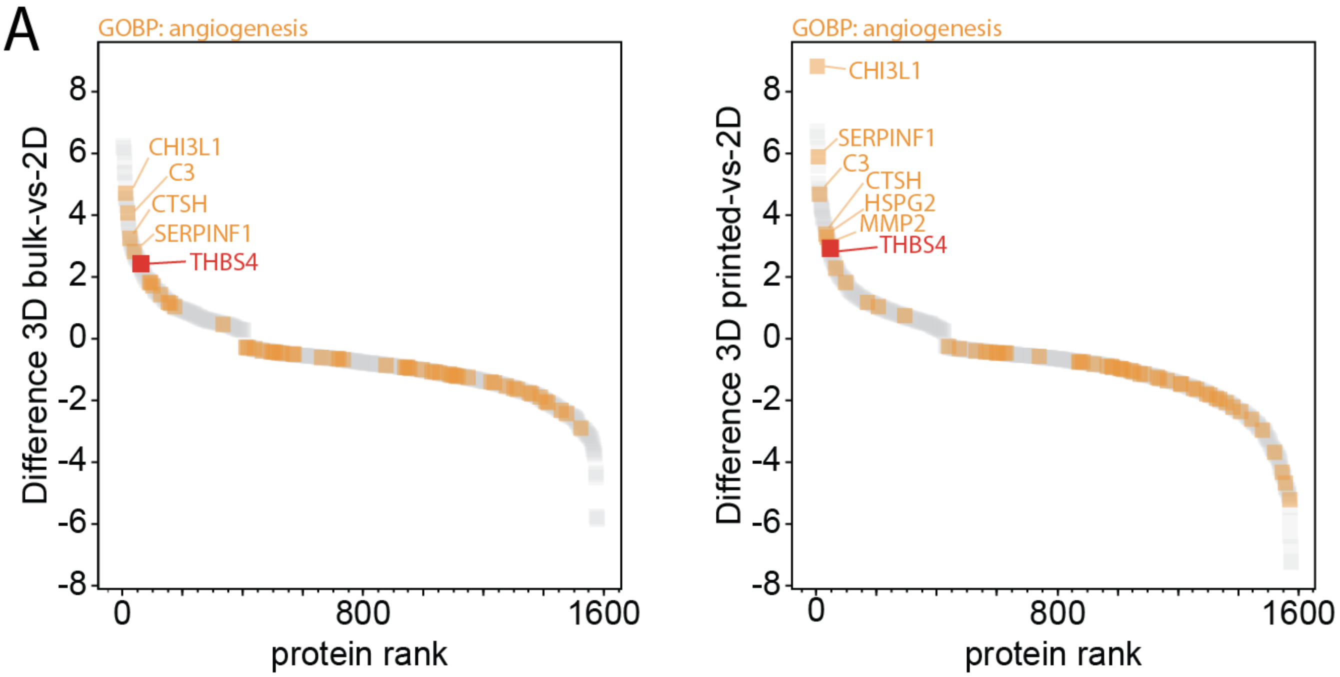


**Supplementary FigureS3**.**THBS4 is secreted in the 3D microenvironment**. (A) Protein rank reporting significant proteins in the 3D-bulk-VS-2D (left plot) and the 3D-printed-VS-2D comparisons (right plot). Terms associated with the GPBP "angiogenesis" are highlighted in orange.
